# Neuronal overexpression of *DYRK1A/minibrain* alters motor decline, neurodegeneration and synaptic plasticity in *Drosophila*

**DOI:** 10.1101/370924

**Authors:** Simon A Lowe, Maria M Usowicz, James JL Hodge

**Author notes:** corresponding authors: Maria Usowicz and James Hodge.

## Abstract

Down syndrome (DS) is characterised by abnormal cognitive and motor development, and later in life by progressive Alzheimer’s disease (AD)-like dementia, neuropathology, declining motor function and shorter life expectancy. It is caused by trisomy of chromosome 21 (Hsa21), but how individual Hsa21 genes contribute to various aspects of the disorder is incompletely understood. Previous work has demonstrated a role for triplication of the Hsa21 gene *DYRK1A* in cognitive and motor deficits, as well as in altered neurogenesis and neurofibrillary degeneration in the DS brain, but its contribution to other DS phenotypes is unclear. Here we demonstrate that overexpression of *minibrain* (*mnb*), the *Drosophila* ortholog of *DYRK1A*, in the *Drosophila* nervous system accelerated age-dependent decline in motor performance and shortened lifespan. Overexpression of *mnb* in the eye was neurotoxic and overexpression in ellipsoid body neurons in the brain caused age-dependent neurodegeneration. At the larval neuromuscular junction, an established model for mammalian central glutamatergic synapses, neuronal *mnb* overexpression enhanced spontaneous vesicular transmitter release. It also slowed recovery from short-term depression of evoked transmitter release induced by high-frequency nerve stimulation and increased the number of boutons in one of the two glutamatergic motor neurons innervating the muscle. These results provide further insight into the roles of *DYRK1A* triplication in abnormal aging and synaptic dysfunction in DS.

**Author summary:** Down syndrome (DS) is caused by three copies of chromosome 21 instead of the usual two. It is characterised by cognitive and motor deficits, which worsen with age resulting in Alzheimer’s disease (AD). Which genes on chromosome 21 cause these phenotypes is incompletely understood. Here we demonstrate that neuronal overexpression of *minibrain*, the *Drosophila* ortholog of the chromosome 21 gene *DYRK1A*, causes age-dependent degeneration of brain neurons, accelerates age-dependent decline in motor performance and shortens lifespan. It also modifies presynaptic structure, enhances spontaneous transmitter release and slows recovery from short-term depression of synaptic transmission at a model glutamatergic synapse. These findings give insight into the role of *DYRK1A* overexpression in aberrant aging and altered information processing in DS and AD.

## Introduction

Down syndrome (DS, also known as Down’s syndrome) or trisomy 21 is caused by the presence of three copies of chromosome 21 (Hsa21) instead of the usual two [1]. It is characterised by cognitive impairment [2] and the delayed and incomplete acquisition of motor skills [3] as a result of abnormal development of the nervous system [4]. Individuals with DS almost invariably develop Alzheimer’s disease (AD)-like symptoms (AD-DS). These include progressive dementia after 40 years of age, the onset of amyloid plaques, neurofibrillary tangles and neurodegeneration after 10 – 20 years [5, 6], faster age-dependent motor decline that is an early marker for the onset of cognitive decline and health deterioration [7, 8], and a shorter mean life expectancy by approximately 28 years [9]. Currently there is no treatment for DS or AD; our understanding of the mechanisms of the disorder is incomplete and this hampers the development of effective therapies.

One of the Hsa21 genes, *DYRK1A* (dual specificity tyrosine-phosphorylation-regulated kinase 1A), is a candidate causative gene for the structural and functional changes that occur in the DS brain, and for the associated cognitive and motor deficits [1, 4]. DYRK1A/Dyrk1a mRNA and protein are expressed throughout the brain in humans and rodents, wherein DYRK1A controls aspects of neuronal development and function [10-12]. DYRK1A/Dyrk1a mRNA and protein expression is increased in DS brain and in the brain of different mouse models of DS [10-13], whether the gene is triplicated as part of a genomic segment, as in Dp1Yey, Ts65Dn, Ts1Cje and Tc1 mice, or alone as in TgDyrk1a and TgDYRK1A mice [1].

DS-associated cognitive and motor deficits are replicated by overexpression of *Dyr1ka* in mice [13-22]. However, the contribution of *DYRK1A* overexpression to the faster age-dependent decline in cognitive and motor function in DS is unclear. It is predicted to play a role as Dyrk1a overexpression intensifies with age in the brain of Ts65Dn and Ts1Cje mouse models of DS [12, 15, 23, 24], it is associated with AD-DS-like histopathological changes in the brain of aged Ts65Dn mice [5, 25] and it enhances APP processing and the formation of hyperphosphorylated Tau aggregates in rat hippocampal progenitor cells *in vitro* [26]. The contribution of *DYRK1A* overexpression to the shorter life expectancy in DS has also not been explored.

Cognitive and motor dysfunction in individuals with DS and in mouse models of DS are associated with changes in synaptic plasticity and with changes in the number and structure of GABAergic and glutamatergic brain neurons and synapses [10, 27, 28]. Such modifications have been linked to *Dyrk1a* overexpression [10, 13, 25, 29], but the effects of *Dyrk1a* overexpression on the basic properties of synaptic function have rarely been explored. In one study, there was no change in the frequency of miniature excitatory synaptic currents (mEPSCs) or the probability of electrically-evoked glutamate release in the prefrontal cortex of TgDyrk1a mice [30]. Nevertheless, since Dyrk1a controls the activity of proteins that regulate endocytosis [31] and *DYRK1A* overexpression slows endocytosis of transmitter vesicles in hippocampal presynaptic membranes from TgDYRK1A mice [32], modulation of transmitter release at other glutamatergic synapses is likely.

To investigate the contribution of *DYRK1A* overexpression in the nervous system to various aspects of DS, we overexpressed *minibrain* (*mnb*), the *Drosophila* ortholog of *DYRK1A* [10], in the *Drosophila* nervous system and implemented well-established assays in larvae and adult flies [33-35]. The assays monitored motor impairment and its development with age, lifespan, age-related neurodegeneration, and synaptic dysfunction. Due to their short lifecycle, *Drosophila* are one of the pre-eminent models for aging and neurodegeneration [36], both aspects of DS that are more difficult to investigate in mice. The *Drosophila* larval neuromuscular junction (NMJ) is a well-established model for mammalian central glutamatergic synapses and is easily accessible to electrophysiology [33]. *Mnb* is expressed presynaptically at larval NMJs and reducing its expression changes motor nerve terminal structure and impairs recycling of transmitter vesicles [37]. Overexpression of the *mnb-F* transcript induces a different change in nerve terminal morphology without apparently affecting basal synaptic transmission [37]. Here we report the effects of neuronal overexpression of *mnb-H*, which encodes the longest mnb splice variant [38], on motor function, the rate of motor decline with age, lifespan, age-related neurodegeneration, presynaptic structure, spontaneous transmitter release and recovery from frequency-dependent depression of electrically-evoked transmitter release.

## Results

### Neuronal overexpression of *mnb* produced motor deficits in larvae, accelerated age-dependent motor decline in adult flies and shortened adult lifespan

The effect of *mnb* overexpression in the nervous system on motor function (specifically the *mnb-H* splice variant) was tested using two assays of fly larval locomotion. *Elav>mnb* larvae, overexpressing *mnb* throughout the nervous system under the control of the *Elav-Gal4* driver [39], did not move as far as control larvae (*Elav/+*) in a free movement assay (Fig. 1A), which measures the ability of larvae to perform rhythmic muscle contractions necessary for gross locomotion [40]. They also took longer to complete a self-righting assay (Fig. 1B), which is a more complex motor task requiring larvae to enact a co-ordinated sequence of movements to right themselves after being rolled onto their backs [41]. To assess the impact of neuronal *mnb* overexpression on age-related decline in locomotor function, the performance of the same cohorts of adult flies was assessed in a negative geotaxis assay at different ages [36]. This showed acceleration in *Elav>mnb* flies of the usual age-related decline in performance (*Elav/+*). There was also evident shortening of the lifespan of *Elav*>*mnb* flies, so that the median lifespan was reduced by almost 50% (*Elav/+*, 73 days; *Elav>mnb*, 38 days; Fig. 1D). These results indicate that neuronal overexpression of *mnb* alone produced a motor deficit and abnormal aging characterised by accelerated age-related locomotor impairment and a shorter lifespan.

**Figure 1.**
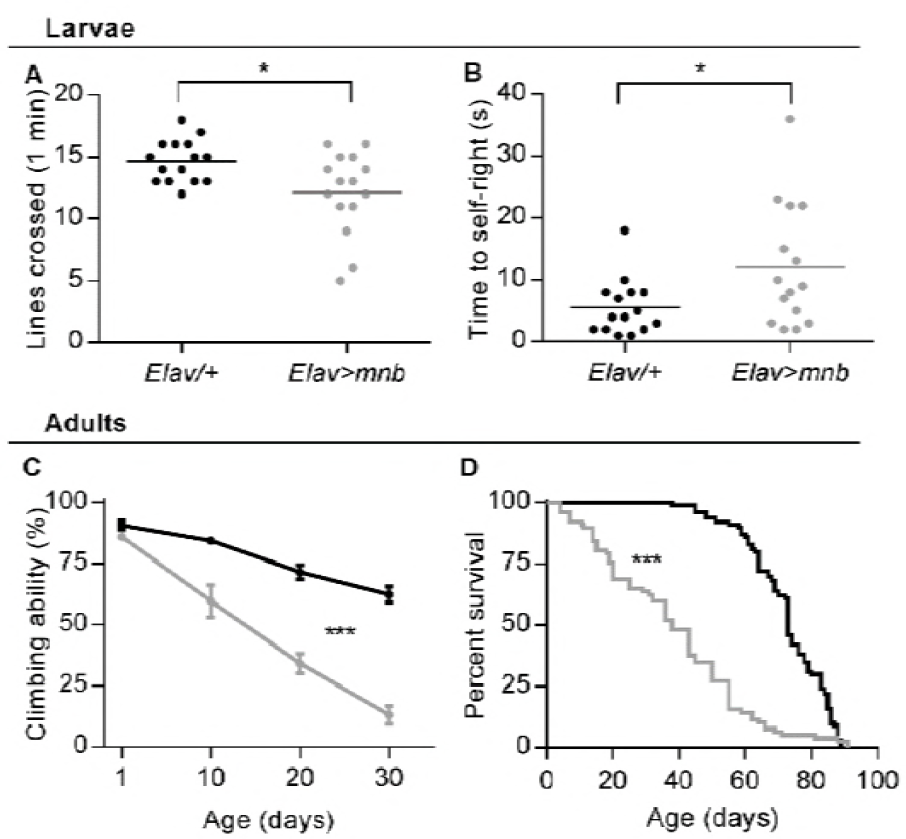
Motor deficits in larvae, accelerated age-dependent motor decline in adult flies and shortened adult lifespan due to neuronal overexpression of *mnb*. (A) *Elav>mnb* larvae crossed fewer lines of a 0.5 cm grid in 60 s than control larvae (*Elav/+*, 14.7 ± 0.44, *n* = 15; *Elav>mnb*, 12.1 ± 0.86, *n* = 15; mean ± SEM, \**P* = 0.014, Student’s *t*-test). (B) *Elav>mnb* larvae took longer than controls to perform a self-righting task (*Elav/+*, 5.5 ± 1.16 s, *n* = 15; *Elav>mnb*, 12 ± 2.56 s; *n* = 15; mean ± SEM, \**P* = 0.029, Student’s *t*-test). Each point in the plots (A, B) is from a different animal; horizontal lines indicate mean values. (C) The age-dependent decline in climbing ability in a negative geotaxis assay was steeper for *Elav>mnb* adult flies than for controls (*F* (3,84) = 13.8; \*\*\**P*< 0.0001, repeated measures two-way ANOVA, *n* = 15 groups of flies); at 1 day old there was no difference in the percentage of flies that climbed successfully (*Elav/+*, 90.59 ± 1.82 %, *n* = 15; *Elav>mnb* ± 85.94 ± 1.46 %, *n* = 15; mean ± SEM, *P* = 0.8262, repeated measures two-way ANOVA and Sidak’s multiple comparison). Values plotted are mean ± SEM, *n* = 15 groups of 10 flies for each genotype. (D) *Elav>mnb* flies had a shorter lifespan relative to controls (*n* = 100 animals per genotype at day 0, \*\*\**P*< 0.0001, log-rank (Mantel-Cox) test).

### Overexpression of *mnb* caused neurodegeneration in adult flies

As *DYRK1A* triplication has been linked to degeneration of brain neurons in AD-DS and in Ts65Dn mice [25, 42], we tested the possibility that neuronal overexpression of *mnb* is sufficient to cause neurotoxicity and age-related neurodegeneration using two established assays of neurodegeneration in adult flies [34, 35]. In the first, *mnb* was overexpressed in the eye through development and adulthood using the *Glass multimer reporter* driver (*GMR-Gal4*) [43]. The *GMR>mnb* flies, but not control flies (*GMR/+*), had a reduced eye surface area and a visible “rough eye” phenotype (Fig. 2A), both of which indicate neural death and the resultant breakdown of the regularly spaced array of ommatidia making up the retina. In a second assay, the *EB1* driver (*EB1-Gal4*) was used to overexpress *mnb* in the ellipsoid body (EB), a subpopulation of neurons within the central complex of the brain implicated in locomotor control (Fig. 2B) [44]. The EB cells also expressed membrane-bound GFP which enabled their visualisation. At 1 day old, there was no difference in the number of GFP-positive EB neurons between control (*EB1/+*) and *EB1>mnb* flies, whereas at day 40 the number of EB neurons was significantly reduced in *EB1>mnb* flies but not in control flies, and the remaining cells were fragmented and misshapen (Fig. 2C). Therefore, neurotoxicity caused by *mnb* overexpression promoted age-related neurodegeneration in a central neuron population.

**Figure 2.**
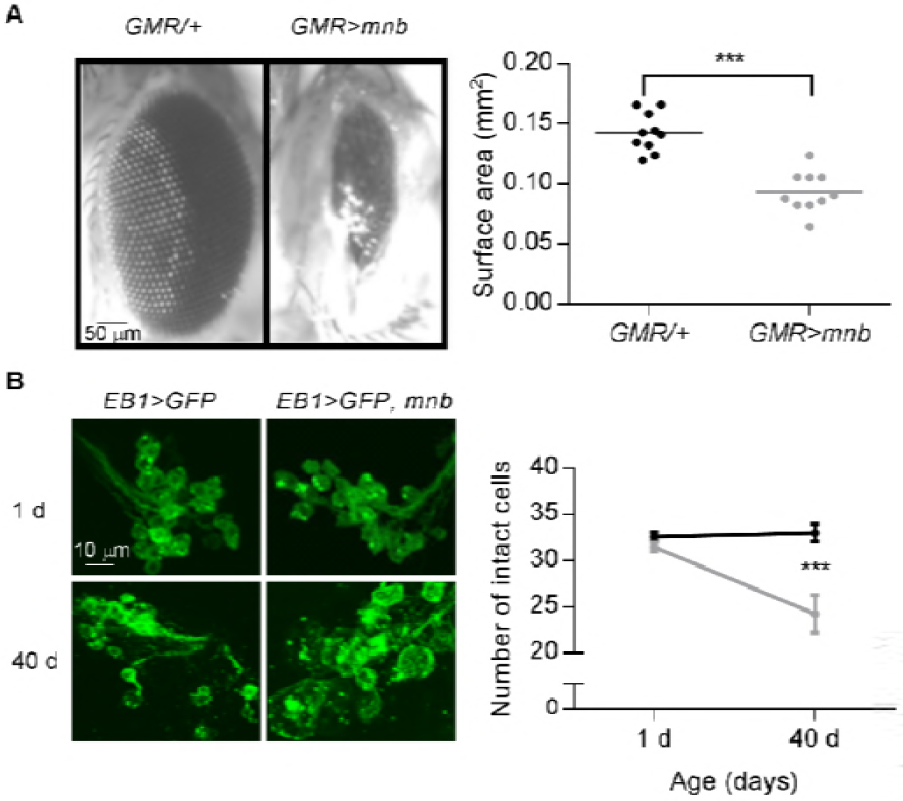
Neurotoxicity and age-related neurodegeneration caused by *mnb* overexpression. (A) (*Left*) Representative images of the eyes of control adult flies (*GMR/+*) and flies with *mnb* overexpression driven in the eye by *GMR-Gal4* (*GMR>mnb*). (*Right*) The surface area of the eyes in *GMR>mnb* flies was reduced (*GMR/+*, 0.14 ± 0.005 mm^2^, *n* = 10; *GMR>mnb*, 0.09 ± 0.005 mm^2^, *n* = 10; mean ± SEM, \*\*\**P* < 0.0001, Student’s *t*-test). Each point in the plot is from a different animal, horizontal lines indicate means. (B) (*Left*) Representative images of clusters of GFP-expressing ellipsoid body neurons in one brain hemisphere, with and without *mnb* overexpression driven by *EB1-Gal4*, in 1 d and 40 d old flies. Calibration bar is 10 μm. (*Right*) The number of intact cells did not differ at 1 d (*EB1>mCD8-GFP*; 32.59 ± 0.48, *n* = 15 (black); *EB1>mCD8-GFP, mnb*; 31.47 ± 0.47, *n* = 15 (grey); mean ± SEM, *P* = 0.757, repeated measures two-way ANOVA and Sidak’s multiple comparison) but decreased between 1 and 40 d in *EB1>mCD8-GFP, mnb* flies; (*F* (1,28) = 10.56; *n* = 15; *P* = 0.003, repeated measures two-way ANOVA). Values plotted are mean ± SEM from 15 flies for each genotype.

### Overexpression of *mnb* in motor neurons increased the number of synaptic boutons at the larval NMJ

To investigate the effect of *mnb* overexpression on presynaptic morphology, *mnb* was overexpressed in glutamatergic motor neurons of *Drosophila* larvae using *OK371-Gal4* [45]. The neuronal membranes were labelled with horseradish peroxidase (HRP) and boutons of the two motor neurons innervating larval muscles 6 and 7 were differentiated according to stronger postsynaptic expression of Discs large (Dlg) opposite 1b (big) boutons than 1s (small) boutons [46]. Analysis of the NMJ in the second abdominal larval segment, A2, showed that *mnb* overexpression affected the morphology of the nerve terminals of only one of the motor neurons; it increased the number of 1b boutons but did not alter the number of 1s boutons (Fig. 3A-B). The effect was not secondary to changes in muscle size, as this did not differ (surface area of muscle 6: *OK371/+*, 44752 ± 1407 μm^2^, *n* = 15; *OK371>mnb*, 44681 ± 3684 μm^2^, *n* = 15, *P* = 0.9857).

**Figure 3.**
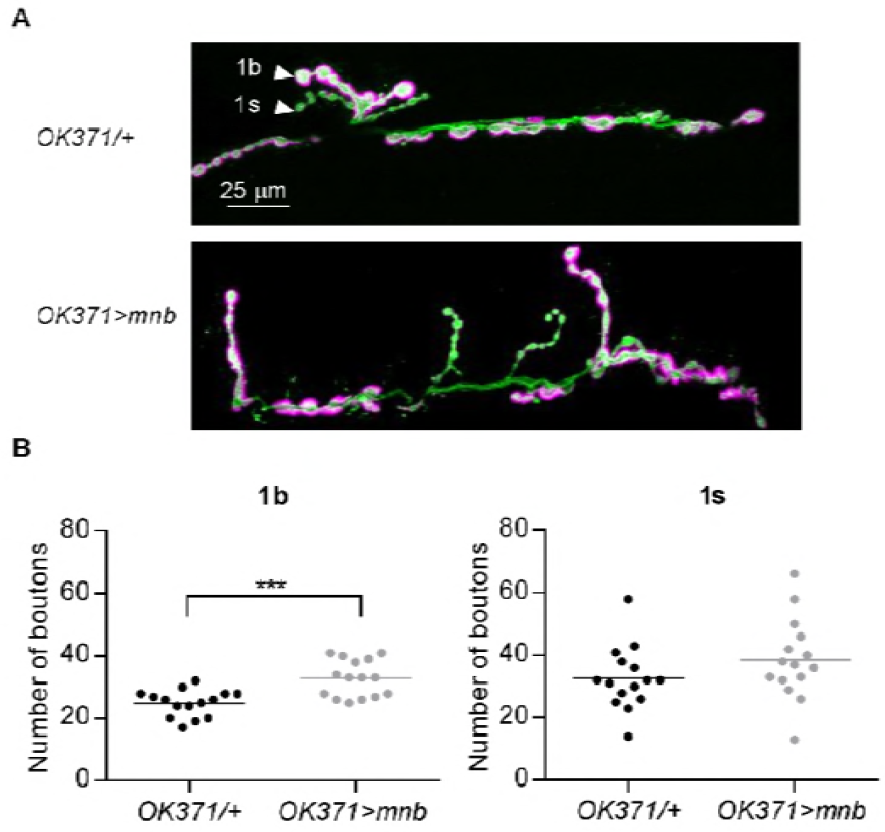
Overexpression of *mnb* in motor neurons increased the number of 1b boutons. (A) Representative images of motor nerve endings at the NMJ of muscle 6/7 in the A2 segment of *OK371/+* (*top*) and *OK371>mnb* (*bottom*) larvae. The neuronal membrane is labelled with HRP (green); type 1b boutons but not type 1s boutons (arrowheads) are preferentially labelled with Dlg (magenta). Scale bar is 25 μm. (B) *OK371>mnb* NMJs had more 1b boutons (*OK371/+*, 24.93 ± 1.11, *n* = 15; *OK371>mnb*, 32.80 ± 1.52, *n* = 15, mean ± SEM, \*\*\**P* = 0.0003, Student’s *t*-test) but there was no difference in the number of 1s boutons (*OK371/+*, 32.6 ± 2.62, *n* = 15; *OK371>mnb*, 38.6 ± 3.34, *n* =15, mean ± SEM, *P* = 0.169, Student’s *t*-test). Each value plotted is from a different animal, horizontal lines indicate means.

### Overexpression of *mnb* in motor neurons altered basal synaptic transmission at the larval NMJ

As neuronal overexpression of *mnb* increased the number of 1b boutons at the larval NMJ, and because previous studies have implicated Dyrk1a/mnb in the control of the recycling of neurotransmitter vesicles [31, 32, 37], we investigated if spontaneous glutamate release was altered by recording spontaneously occurring miniature excitatory junction potentials (mEJPs) with intracellular microelectrodes. Since the muscle 6/7 NMJ is innervated by 1b and 1s boutons, mEJPs usually result from the release of neurotransmitter vesicles from both types of boutons [47]. Our recordings revealed that mEJPs occurred more frequently but were smaller in *OK371>mnb* larvae (Fig. 4A-B). The decrease in amplitude was not secondary to a change in the electrical properties of the muscle as there was no difference in input resistance (*OK371/+*, 3.37 ± 0.69 MΩ, *n* = 8; *OK371>mnb* = 3.99 ± 0.99 MΩ, *n* = 8, *P* = 0.613) or resting potential (*OK371/+*, −69.6 ± 1.48 mV, *n* = 8; *OK371>mnb*, − 67.6 ± 1.55 mV, *n* = 8, *P* = 0.365). Although we did not directly investigate the source of the more frequent smaller mEJPs, the notion that they are due to the observed selective increase in the number of 1b boutons is suggested by the fact that 1b-dependent mEJPs are smaller than 1s-dependent mEJPS [47]. In parallel with the changes in spontaneous synaptic events, there was a small (~11 %) decrease in the mean amplitude of electrically-evoked excitatory junction potentials (EJPs) caused by single stimuli applied to the nerve at a low frequency (0.1 Hz) (Fig. 4C). There was no difference between EJPs in mean rise time (*OK371/+*, 2.67 ± 0.178 ms, *n* = 8; *OK371>mnb*, 3.23 ± 0.33 ms, *n* = 8, *P* = 0.728) or mean time constant of decay (*OK371/+*, 44.9 ± 3.1 ms, *n* = 8; *OK371>mnb*, 36.6 ± 4.6 ms, *n* = 8, *P* = 0.154). The relatively small fall in EJP amplitude is likely to reflect the smaller size of the 1b-dependent component of the EJP relative to that of the 1s-dependent component [47].

**Figure 4.**
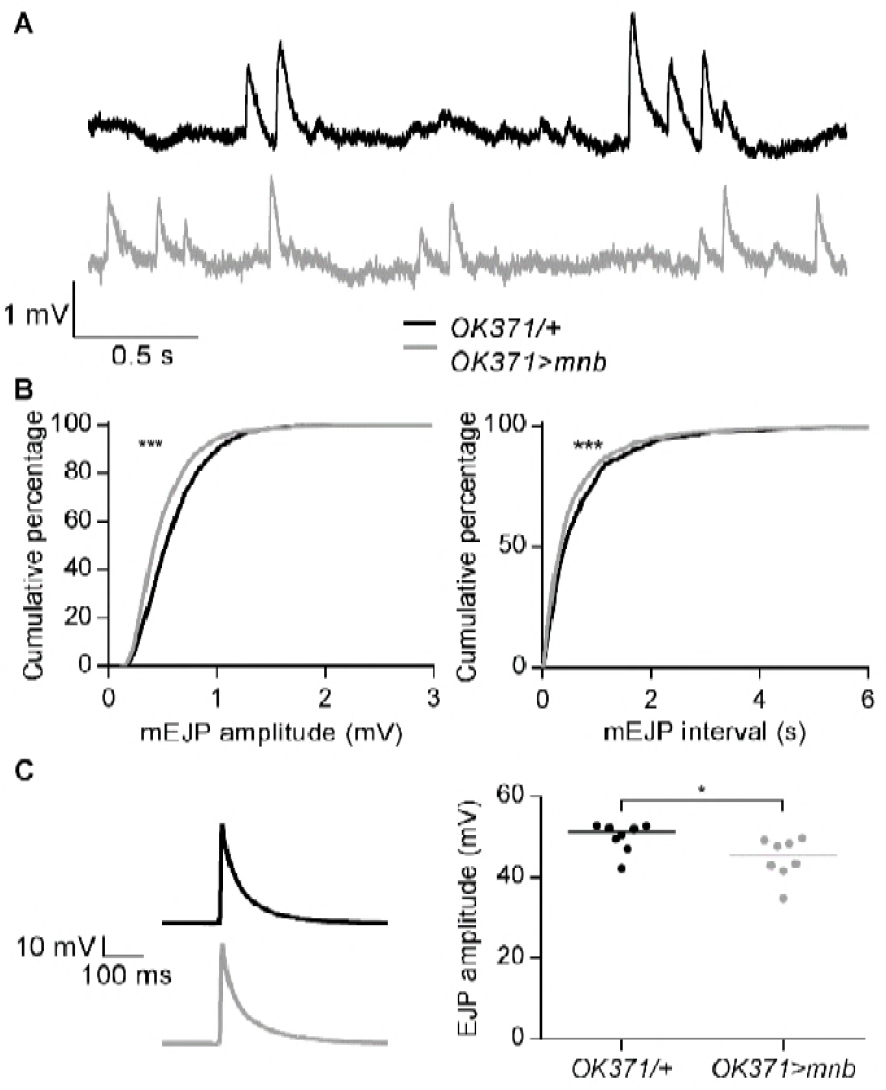
Overexpression of *mnb* in motor neurons altered basal synaptic transmission. (A) Representative voltage recordings (3 s traces) from NMJs of *OK371/+* (black) and *OK371>mnb* (grey) larvae showing spontaneous mEJPs recorded at a membrane potential of −70 mV. (B) Cumulative frequency distributions of mEJP amplitude (*left*) and inter-mEJP interval (*right*) (1600 events, 200 from each of 8 NMJ recordings). *OK371>mnb* mEJPs were smaller (\*\*\**P* < 0.0001) and more frequent (\*\*\**P* < 0.0001), Kolmogorov-Smirnov test. (C) (*Left*) Representative image of a single electrically-evoked EJP at an *OK371/+* NMJ and an *OK371>mnb* NMJ at a membrane potential of −70 mV. (*Right*) The median EJP amplitude (horizontal line) was reduced in *OK371>mnb* NMJs (*OK371/+*, 51.26 mV, *n* = 8; *OK371>mnb*, 45.55 mV, *n* = 8; mean ± SEM, \**P* = 0.0281). Each point plotted is from a different animal.

### Overexpression of *mnb* in motor neurons slowed recovery from frequency-dependent depression at the larval NMJ

To investigate the effects of neuronal *mnb* overexpression on recycling of synaptic vesicles during electrically-evoked transmitter release, EJPs were evoked with pairs of electrical stimuli separated by intervals of varying duration (10 ms – 10 s) or with repeated trains of 10 stimuli applied at a high frequency (10 Hz, a frequency 100 times higher than that at which the single EJPs were evoked) [48]. At control NMJs, paired pulses separated by intervals shorter than 200 ms caused depression of the amplitude of the second EJP relative to that of the first and the depression was stronger for shorter inter-stimulus intervals (Fig. 5A). The dependence of paired-pulse depression on interval duration was unaltered in *OK371>mnb* larvae (Fig. 5A), indicating that *mnb* overexpression did not alter release from a readily releasable pool of vesicles [49]. When transmitter release was evoked at control NMJs with a train of 10 stimuli at 10 Hz, there was rapid depression of the EJP amplitude by ~20% within the first 3 events (Fig. 5B). In the one-minute interval before the next train, the EJP amplitude recovered fully so that the amplitude of the first EJP in the second train was the same as in the first train (Fig. 5B). This ability to recover did not wane during the recording; the amplitude of the first EJP in each train did not differ between 8 trains (Fig. 5B). These effects are consistent with previous studies [48] and confirm rapid depletion and replenishment of the readily releasable pool of vesicles [49]. However, the same pattern of nerve stimulation produced different effects at *OK371>mnb* NMJs (Fig. 5B). The percentage decrease in EJP amplitude during each train was the same as at control NMJs, but the depression was not fully reversed during the intervals between trains, so that the first EJP in each train was smaller than the first EJP in the preceding train. The depression in amplitude accumulated over the 8 trains, resulting in an overall fall of 10%. To confirm that the changes in EJP amplitude were due to presynaptic changes in transmitter release and were not postsynaptically mediated by a decrease in the unitary depolarisations comprising each EJP, we measured the amplitudes of 200 mEJPs immediately before and 200 mEJPs immediately after the series of trains at each NMJ. At both control and *OK371>mnb* NMJs, the cumulative distribution of mEJP amplitudes before and after a series of trains was similar; although they were not identical, the observed slight increase in the number of larger mEJPs cannot explain the decline in EJP amplitude (Fig. 5C). These results show that *mnb* overexpression slowed replenishment of the readily releasable pool of vesicles, an effect consistent with the reported slowing of endocytosis of transmitter vesicles by *DYRK1A* overexpression [32].

**Figure 5.**
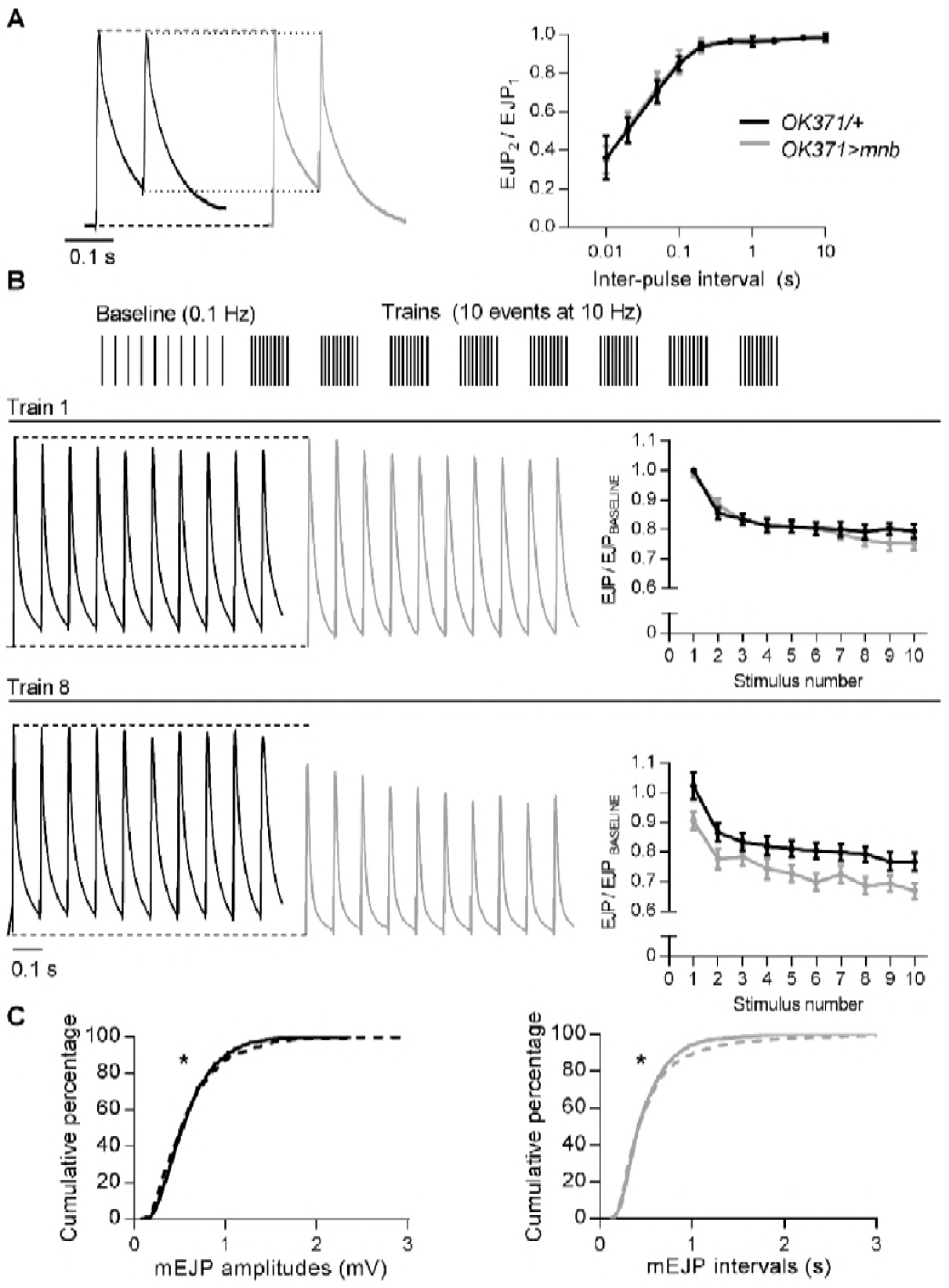
*mnb* overexpression in motor neurons slowed recovery from frequency-induced depression. (A) *(Left)* Representative pairs of stimulus evoked EJPs at *OK371/+* (black) and *OK371>mnb* (grey) NMJs. Dashed lines compare the first EJP (EJP_1_) and dotted lines compare the second EJP (EJP_2_). (*Right*) Plots of paired-pulse ratio (EJP_2_/EJP_1_, mean ± SEM, *n* = 8 for each genotype) against inter-pulse interval reveal no difference in synaptic depression (*F* (9, 126) = 0.1343; *n* = 8; *P* = 0.9987, repeated measures two-way ANOVA). (B) High-frequency stimulation protocol. 10 EJPs were evoked at 0.1 Hz to establish a mean baseline amplitude followed by 8 trains of 10 EJPs at 10 Hz, at one minute intervals. *(Left)* Representative traces of train 1 (*upper*) and train 8 (*lower*) recorded from *OK371/+* and *OK371>mnb* NMJs. Dashed lines compare EJP_1_ in each train. *(Right)* Plots of EJP amplitude during trains 1 and 8, expressed as a fraction of baseline amplitude (mean ± SEM, *n* = 8 for each genotype). During the first train, the decline in EJP amplitude was unaffected by genotype (*F* (1, 14) = 0.22, *P* = 0.6486, repeated measures two-way ANOVA). In the eighth train, EJPs were smaller at *OK371>mnb* NMJs (*F* (1,14) = 4.98, *P* = 0.0426) but the rate of decline during the 8^th^ train was not different (*F* (9, 126) = 0.99, *P* = 0.4540, repeated measures two-way ANOVA). (C) Cumulative frequency distributions of mEJP amplitudes (*n* = 1600 from 8 NMJ recordings) from immediately before (solid line) and immediately after (dashed line) the trains, for *OK371/+* (*left*) and *OK371>mnb* (*right*) NMJs; after the trains, a minority of mEJPs were larger and occurred less frequently in both *OK371/+* (\**P* = 0.0117) and *OK371>mnb* (\**P* = 0.018) NMJs (log-rank (Mantel-Cox) test).

## Discussion

This study demonstrated that neuronal overexpression of *mnb*, the *Drosophila* ortholog of DYRK1A, is sufficient to induce motor impairment, accelerate age-related decline in motor performance, shorten lifespan and cause age-dependent neurodegeneration. This study also found that neuronal *mnb* overexpression at a glutamatergic synapse alters presynaptic structure, modifies basal synaptic transmission and delays recovery from short-term synaptic depression.

People with DS have impaired motor skills which are evident from childhood and are caused by abnormal development of the nervous system [3, 4]. Later, in middle age, they undergo faster age-dependent motor decline, which is an early marker of future dementia, comorbidities and mortality, and is likely caused by histopathological changes in the brain [7, 8]. The life expectancy of people with DS is about 28 years shorter than the general population [9]. By taking advantage of the relatively short life cycle of *Drosophila* and driving overexpression of *mnb* in neurons, we have demonstrated a potential role for neuronal DYRK1A overexpression in the accelerated age-dependent decline of motor function and shortening of life expectancy in DS. The genetic basis of these aspects of DS is more difficult and costly to explore in mouse models of DS, due to the longer time required to study aged mice. Our finding that *mnb* overexpression causes age-related neurodegeneration confirms previous studies inferring a link between *DYRK1A* overexpression and degeneration and loss of neurons [10, 15, 25, 42], which is associated with faster age-related decline in motor and cognitive function in DS and AD-DS. Our results also reinforce the conclusion from earlier studies with adult mice overexpressing *DYRK1A or Dyrk1a*, alone or as part of a chromosomal segment, that triplication of *DYRK1A* is likely to contribute to motor deficits in DS [13-19].

In addition to the smaller brain size and fewer brain neurons in DS and mouse models of DS, there are alterations in the structure of brain synapses that are predicted to modify synaptic function [4, 28, 50]. A previous study showed that DYRK1A overexpression in mice changes postsynaptic morphology in the cortex and in cultured cortical neurons by reducing the number and length of dendrites and by reducing the number of dendritic spines but elongating their shape [51]. It also decreased the number of synapses formed. Our study shows that *mnb* overexpression changes presynaptic structure and that this happens in a neuron-specific manner; *mnb* overexpression in the two glutamatergic motoneurons innervating the larval NMJ increased the number of 1b boutons without changing the number of 1s boutons. These data are consistent with a previous study which demonstrated that reduced levels of *mnb* caused a decrease, and increased levels of the *mnb-F* transcript an increase, in the number of boutons at the NMJ [37], but did not differentiate between 1b and 1s boutons.

The cognitive and motor deficits in DS arise from aberrant information processing in the brain that is likely due, in part, to changes in synaptic transmission or synaptic plasticity. Individuals with DS have impaired synaptic plasticity in the motor cortex [27]. Our finding that *mnb* overexpression slows replenishment of the readily releasable pool of vesicles, and also modifies basal synaptic transmission, confirms a previous suggestion that DYRK1A overexpression contributes to synaptic dysfunction and cognitive deficits associated with DS, made on the basis of the observed slowing of endocytosis of transmitter vesicles in cultured mouse hippocampal neurons overexpressing human *DYRK1A* [32]. The effects of *DYRK1A* on synaptic function may be splice variant specific as we found that overexpression of the *mnb-H* transcript caused a decrease in mEJP and EJP amplitude, whereas overexpression of *mnb-F* in a previous study caused an increase in mEJP amplitude and no change in EJP amplitude at the larval NMJ [37].

The effects of neuronal *mnb* overexpression on larval NMJ function replicate some, but not all, the documented changes in glutamatergic synaptic transmission in the brain of mouse models of DS. These include a decrease in the amplitude of spontaneous excitatory postsynaptic currents (sEPSCs) in neocortical neurons of Ts65Dn mice [52], compromised glutamate release in response to stimuli trains at hippocampal CA1 synapses of Ts1Cje mice [53] and a decrease in EPSC amplitude in hippocampal CA3 neurons of Ts65Dn mice [54]. However, in contrast to the increase in mEJP frequency caused by *mnb* overexpression at the larval NMJ, electrophysiological studies have found a decrease in the frequency of mEPSCs in hippocampal CA3 neurons of Ts65Dn mice, sEPSCs in neocortical neurons of Ts65Dn mice and sEPSCs in neurons derived from trisomy 21 induced pluripotent stem cells, or no change in mEPSC frequency in the prefrontal cortex of TgDyrk1a mice or mossy fibre-CA3 synapses in Tc1 mice [28].

Our study further elucidates the role of DYRK1A triplication in various DS phenotypes. It supports the future development of pharmacological inhibitors of DYRK1A as treatments for multiple aspects of DS and AD [10, 12]. Further work is necessary to fully understand interactions between DYRK1A and other triplicated Hsa21 genes in DS, in specific cell types and during defined periods of development and ageing.

## Materials and Methods

### Animals

Flies were raised with a 12 h:12 h light dark cycle with lights on at ZT 0 (Zeitgeber time) on standard *Drosophila* medium (0.7% agar, 1.0% soya flour, 8.0% polenta/maize, 1.8% yeast, 8.0% malt extract, 4.0% molasses, 0.8% propionic acid, 2.3% nipagen) at 25°C. Flies were transferred to vials containing fresh medium twice weekly. The *OK371-Gal4* (Bloomington stock center numbers: 26160)*, Elav-Gal4* (87060)*, GMR-Gal4* (9146) flies were obtained from the Bloomington *Drosophila* Stock Centre. *Canton Special white-* (*CSw-*) flies were a gift from Dr S. Waddell (University of Oxford), *UAS-mnb* flies (*minibrain-H*, CG42273) were kindly provided by Dr Kweon Yu (Korea Research Institute of Bioscience and Biotechnology), *EB1-Gal4; UAS*-*mCD8-GFP* flies were donated by Dr F. Hirth (Kings College London).

### Behaviour

*mnb* expression was driven throughout the nervous system using *Elav-Gal4* [39] for experiments investigating behaviour of wandering third instar larvae, the number of boutons at the larval NMJ, and synaptic transmission at the larval NMJ. All behavioural experiments took place at 25°C. Larval locomotor experiments were conducted on a 9.5 cm petri dish containing 1.6% agarose. A single third instar wandering larva was selected, washed in a drop of distilled H_2_O, transferred to the agarose and allowed 30 s to acclimatise. To analyse free movement, the dish was placed over a 0.5 cm grid and the number of lines the larva crawled across in one minute was counted by eye. The self-righting assay was conducted as described elsewhere [41, 55]; the larva was gently rolled onto its back on the agarose using a fine moistened paintbrush, held for one second and released, and the time for it to right itself was recorded.

The negative geotaxis assay was performed as described previously [56]. A cohort of 10 flies was transferred without anaesthesia to an empty 9.5 cm tube with a line drawn 2 cm from the top. After 1 minute acclimatisation, the vial was sharply tapped 3 times to knock the flies to bottom. The number of flies to climb past the line within 10 s was recorded. 15 cohorts of 10 flies were tested for each genotype. Age-dependent changes in climbing were assessed by repeating the negative geotaxis assay at 10, 20 and 30 days post-eclosure [57]. For the survival assay, 10 cohorts of 10 once-mated females were transferred to a vial of fresh food twice weekly and the number of surviving flies recorded at each transfer.

### Antibody staining and visualisation at the NMJ

Wandering third instar larvae were dissected in ice-cold, Ca^2+^-free HL3.1-like solution (in mM: 70 NaCl, 5 KCl, 10 NaHCO_3_, 115 sucrose, 5 trehalose, 5 HEPES, 10 MgCl_2_) to produce a larval “fillet” [58]. The fillet was fixed for 30 min in 4% paraformaldehyde (Sigma Labs), washed three times in 1% Triton-X (Sigma Labs) and blocked for one hour in 5% normal goat serum (Fitzgerald Industries) and 1% Triton-X at room temperature. It was incubated overnight in 1/500 FITC-conjugated anti-horseradish peroxidase (HRP-FITC) (Jackson Immunoresearch Laboratories) and 1/500 mouse anti Discs large (Dlg) primary antibody, then for two hours in 1/500 AlexaFluor 633-conjugated goat anti-mouse secondary antibody at room temperature. Each fillet was washed and mounted on a coverslip in Vectashield (Vector Laboratories). Z-series of NMJs were imaged on a Leica SP5-II confocal laser-scanning microscope using an oil immersion 40 × objective. The number of boutons at the NMJ of muscle 6/7 in segment A2 was counted manually. ImageJ (rsb.info.nih.gov/ij/) was used to manually outline muscle 6 and hence calculate their area.

### Neurotoxicity

Overexpression of *mnb* was driven in the eye using the *Glass multimer reporter* (*GMR-Gal4*). Images of the whole head of 1-2 day old flies were taken via a Zeiss AxioCam MRm camera attached to a stereomicroscope (Zeiss SteREO Discovery.V8, up to 8× magnification), and the surface area of the eye was calculated by manually outlining the eye in ImageJ (rsb.info.nih.gov/ij/). Overexpression of *mnb* in GFP-tagged ellipsoid body (EB) ring neurons was achieved by crossing *EB1-Gal4; UAS*-*mCD8-GFP* flies with *UAS*-*mnb* flies. Following published methods [59], adult brains were dissected, fixed for 30 minutes in 4% paraformaldehyde and mounted on a coverslip in Vectashield (Vector Laboratories). Slides were imaged on a Leica SP5-II confocal laser scanning microscope using an oil immersion 40 × objective. A Z-stack of 25 images at 1 μm increments was captured and combined into a 3-D projection using ImageJ (rsb.info.nih.gov/ij/); analysis was performed by scrolling through all 25 images and counting the number of intact cells in one brain hemisphere.

### Electrophysiology

Wandering third instar larvae were dissected as for antibody staining. The motor nerves were severed just below the ventral ganglion and the brain was removed. CaCl_2_ (1 mM) was added to the bath solution for intracellular recording from muscle 6 of abdominal segments 2-4. Most recordings were made in the presence of thapsigargin (2 μM), which minimises contraction and hence facilitates intracellular recording, without affecting amplitudes of mEJPS or EJPs [47, 60]. Sharp microelectrodes (thick-walled borosilicate capillaries, pulled on a Sutter Flaming/Brown P-97 micropipette puller) were filled with 3M KCl and had resistances of 20 – 30 MΩ. For recording of stimulus evoked excitatory junction potentials (EJPs), severed nerves were drawn into a thin-walled glass-stimulating pipette and stimulated with square-wave voltage pulses (0.1 ms, 10 V, A-M Systems Model 2100 Isolated Pulse Simulator), 10 times at 0.1 Hz. EJPs and spontaneously-occurring miniature EJPs (mEJPs) were recorded at a controlled room temperature of 22-25°C with a Geneclamp 500 amplifier (Axon Instruments) and were further amplified with a LHBF-48x amplifier (NPI Electronic). The membrane potential was allowed to stabilise for one minute, the initial value was recorded, and then set to −70 mV by injecting current with the Geneclamp 500 amplifier. The muscle input resistance was measured by injecting current using the Axon Geneclamp 500, to bring the membrane potential to −100, −80, −60 and −40 mV, and subtracting the electrode resistance from the slope of the resulting voltage/current graph. Voltage signals were low-pass filtered at 1.67 kHz (10 kHz 4 pole Bessel on Geneclamp 500, 1.7 kHz 8-pole Bessel on LHBF-48x) and digitised at 25 kHz by a CED-1401 plus A/D interface (Cambridge Electronic Design, UK) using Spike2 software (v. 5.13) (CED, Cambridge, UK). Recordings were discarded if the initial resting membrane potential was more positive than −60 mV or varied by more than 10% throughout the recording. Synaptic potentials were analysed off line using Strathclyde Electrophysiology Software WinEDR (v3.5.2) and GraphPad Prism (v.6). All synaptic events were verified manually.

Amplitudes and intervals of mEJPs were compared by creating a cumulative distribution for each genotype of 1600 measurements across 8 animals, with each animal contributing 200 values. To analyse the mEJP waveform, a mean mEJP was constructed for each recording from events showing a single clear peak and a smooth decay, so as to prevent distortion of the waveform by closely occurring mEJPs. A single exponential was fitted to the decay of the mean mEJP and the 10-90% rise-time was measured. Time zero for the exponential fit was set to the time at the peak of the mEJP. EJPs were analysed by forming a mean of 10 events, measuring the 10-90% rise-time of the mean event, and fitting the decay with the sum of three exponentials (time zero was set at the time of the peak). A mean weighted time constant of decay was calculated as A_1_.τ_1_ + A_2_.τ_2_ + A_3_.τ_3_, where A_1_, A_2_ and A_3_ are the fractional amplitudes of the three components, and τ_1_, τ_2_ and τ_3_ are their time constants.

For paired pulse analysis, two EJPs were evoked, separated with intervals varying in duration between 0.01 and 10 s. The amplitude of the second event was calculated as a fraction of the amplitude of the first. Pairs of stimuli were separated by 30 s. For high–frequency stimulation, trains of 10 events evoked at 10 Hz were repeated 8 times at 1 min intervals. The amplitude of each event was expressed as a fraction of a baseline value, defined as the mean amplitude of 10 single EJPs evoked at 0.1 Hz. To compare mEJP amplitude before and after the trains, 200 mEJP amplitudes were measured immediately before the first train, and another 200 immediately after the eighth train. The measured values were pooled from 8 NMJs for each genotype and cumulative amplitude distributions compared.

### Statistical analysis

Statistical analysis was conducted in GraphPad Prism (v. 6, La Jolla, CA). Data were tested for normality using the Kolmogorov-Smirnov test; where appropriate means were compared using Student’s unpaired *t*-test, or medians were compared using a Mann-Whitney *U* test. EJPs evoked by pairs or trains of stimuli were compared using repeated measures 2-way ANOVA. Cumulative distributions were compared with a Kolmogorov-Smirnov test. Survival curves were compared with a Mantel-Cox test. Data are given as median or mean ± SEM. *n* is given per genotype. An α level of *P* < 0.05 was considered significant.

